# Genome-wide identification and expression analysis of the invertase gene family in common wheat (*Triticum aestivum* L.)

**DOI:** 10.1101/2021.12.29.474404

**Authors:** Chao Wang, Guanghao Wang, Xiaojian Qu, Xiangyu Zhang, Pingchuan Deng, Wanquan Ji, Hong Zhang

## Abstract

**Background:** The degradation of sucrose plays an important role in the process of crop biomass allocation and yield formation. Invertase (INV) irreversibly catalyzes the conversion of sucrose into glucose and fructose, which doomed its’ important role in plant development and stress tolerance. However, the functions of INV genes in wheat, one of the most important crops, were less studied due to the polyploidy.

**Results:** Here, we systematically analyzed the INV gene family based on the latest published wheat reference genomic information. A total of 126 *TaINV* genes were identified and classified into three classes based on the phylogenetic relationship and their gene structure. Of which, 11 and 83 gene pairs were identified as tandem and segmental duplication genes respectively, while the Ka/Ks ratios of tandem and segmental duplication *TaINV* genes were less than 1. Expression profile analysis shows that 18 *TaINV* genes have tissue-specific expression, and 54 *TaINV* genes were involved in stress response. Furthermore, RNA-seq showed that 35 genes are differentially expressed in grain weight NILs N0910-81L/N0910-81S, in which 9 *TaINVs* were stably detected by qRT-PCR at three time-points, 4, 7 and 10 DPA. Four of them (*TaCWI47*, *TaCWI48*, *TaCWI50* and *TaVI27*) different expressed between the NILs resided in 4 QTL segments (*QTGW.nwafu-5DL.1*, *QTGW.nwafu-5DL.2*, *QTGW.nwafu-7AS*.1 and *QTGW.nwafu-7AS.2*). These findings facilitate function investigations of the wheat INV gene family and provide new insights into the grain development mechanism in wheat.

**Conclusions:** Our results showed that allopolyploid events were the main reason for the expansion of the *TaINV* gene family in hexaploid wheat, and duplication genes might undergo purifying selection. The expression profiling of *TaINV* genes implied that they are likely to play an important role in wheat growth and development and adaption to stressful environments. And *TaCWI47*, *TaCWI48*, *TaCWI50* and *TaVI27* may have more important roles in grain developments. Our study lay a base for further dissecting the functional characterization of *TaINV* family members.

## Introduction

Since the separation of invertase (ROBERT 1976; Roberts 1973), the structure and function of invertase have gradually been discovered. Invertase irreversibly catalyzes the conversion of sucrose into glucose and fructose (Arnd Sturma 1999). Based on their optimum pH, the invertase isoenzyme forms can be classified into two subfamilies, neutral/alkaline invertases (A/N-Invs) and acid invertases (Ac-Invs) (Koch 2004). Ac-Invs is also called β-D-fructofuranosidase (EC3.2.1.26) because it can hydrolyze other oligosaccharides containing Fru such as raffinose and stachiose (Lammens et al. 2009). And Ac-Invs are glycosylated proteins classified in glycoside hydrolase family 32 (GH32). According to their localization in apoplast and vacuole, Ac-Invs can be be further divided into cell wall invertase (CWI) and vacuolar invertase (VI) (Sherson SM 2003). Meanwhile, A/N-Invs are nonglycosylated proteins classified in glycoside hydrolase family 100 (GH100) and specifically hydrolyze sucrose. A/N-Invs exist as two different isoforms with optimum pH either close to 6.5 or 8, named N-Inv or A-Inv, respectively. However, A/N-Invs is better known as cytoplasm invertases (CI), due to its localization in the cytoplasm (Vargas & Salerno 2010).

The first cDNA cloning and expression of CWI in carrots marked the molecular level of invertase research (Chrispeels 1990). It is believed that Ac-INV originating from respiratory eukaryotes and aerobic bacteria (Ji et al. 2005) and CI seem to be arised from cyanobacteria (Arnd Sturma 1999). The evolutionary relationship between cell wall invertase and vacuolar invertase is still controversial (Ji et al. 2005; Van den Ende 2013). Ji et al. (2005) believe that vacuolar invertase evolved from cell wall invertase, while Van et al. (2013) hold the opposite view. They thought higher plant CWIs evolved from an ancestor shared with VIs from lower plant species. Ac-Invs contains three conserved motifs of NDPNG (β-fructofuranosidases motif), RDP, and WECP(V)D, which are necessary for the catalytic activity (Ji et al. 2005). Moreover, three amino acids (DPN) of the β-fructofuranosidases motif is the smallest exon known to plant species and skipped in cold stressed potato (Bourney et al. 1996). In contrast to CWI, VI contain a C-terminal extension (Sturm 1999) and a much longer and complex N-terminal propeptide region (NTPP), containing a dileucine motif, an YXXΦ (X is any amino acid, Φ is a hydrophobic residue) motif, a basic region motif and a transmembrane domain (TMD) (Ji et al. 2005; Xiang & Van den Ende 2013).

Numerous studies have shown that INV plays multiple roles in plants, including development, sucrose metabolism, and stress responses (Morey et al. 2018; Su et al. 2018; Wan et al. 2018). CI is a critical player in the regulation of plant roots and their reproductive development (Lou et al. 2007; Martin et al. 2013; Takanori Maruta 2010). For example, mutation of CYT-INV1 in arabidopsis and rice resulted in a shorter primary root (Jia et al. 2008; Qi et al. 2007), accompanied by a smaller leaves and siliques in arabidopsis (Qi et al. 2007) and delayed flowering and partial sterility in rice (Jia et al. 2008). CWI has a role in determining of grain size, grain number and grain quality. The decrease of CWI activity inhibits the grain development, resulting in yields decrease in maize (McLaughlin & Boyer 2004), rice (Cho et al. 2005; Tatsuro Hirose 2002; Wang et al. 2008), tomato (Zanor et al. 2009) and wheat (Sonia Dorion 1996). Similarly, mutation of the CWI gene *INCW2* in maize reduced maize grain weight by 80% (Chourey 1992). Conversely, inhibiting the expression of the CWI inhibitor gene in tomato (*INVINH1*) and soybean (*GmCIF1*) can improve fruit and seed set (Jin et al. 2009; Qin et al. 2016; Tang et al. 2017). Over-expression of CWI enhanced grain filling and yield in rice and maize (Li et al. 2013; Tatsuro Hirose 2002; Wang et al. 2008). VI has the roles in the determination of hexose-to-sucrose ratio (Ellen M. Klann 1996; Rita Zrenner 1996), cell elongation (Sergeeva et al. 2006; Wang et al. 2010), and cell expansion (Morey et al. 2018). The reduction of vacuolar invertase activity caused a decline in photosynthesis and decreased carbon export-related metabolic pathways and sink organs (Nagele et al. 2010).

The identification of 17 invertase genes in *Arabidopsis thaliana* and 18 invertase genes in *Oryza sativa* (Cho et al. 2005; Ji et al. 2005; Najat Haouazine-Takvorian 1997), laid the foundation for the subsequent research on gene families of other plants. Subsequently, 21 invertase genes in *Zea may*, 24 in *Populus*, 18 in *Malus domestica*, 32 in *Glycine max*, 19 in *Sorghum bicolor*, and 19 in *Brachypodium distachyon* were identified by genome-wide analyses (Chen et al. 2015; Juarez-Colunga et al. 2018; Su et al. 2018; Tae Kyung Hyun 2011; Wang et al. 2017). Wheat (*Triticum aestivum* L.) is one of the most important cultivated crops. According to this ratio, wheat encodes more than a hundred INVs in this hexaploid plant with an estimated 107,891 high-confidence genes (Consortium 2018). To date, only eight *TaCWI* genes (*IVR1.1–1A*, *IVR1.2–1A*, *IVR1.1–3B*, *IVR1–3A*, *IVR1–4A*, *IVR1–5B*, *IVR1.2–3B* and *IVR1-5D*). one *TaCI* gene (*Ta-A-Inv*) and four *TaVI* genes (*TaVI1*, *TaVI2*, *TaVI3* and *Ivr5*) were reported in wheat (Koonjul et al. 2005; Liu et al. 2015; Webster et al. 2012). *Ta-A-Inv* is up-regulated in wheat leaves after osmotic stress, low-temperature treatment (Vargas et al., 2007), and *Puccinia striiformis* f. sp. tritici infection (Liu et al., 2015). To understand the landscape of wheat INVs and the molecular mechanisms determining grain size, we used whole-genome scanning to identify INV family members.

Herein, a total of 126 INV genes were identified in wheat. They were categorized into 3 classes and their characteristics of conserved motif, gene structure, and gene duplication for different classes were analyzed. We also exhibited here the phylogenetic relationship and the syntenic correlation among wheat, rice, maize, arabidopsis, and soybean. Expression profiling implied the possible roles in regulating development and responding to abiotic stresses. Therefore, this study provides a cue for further research on the functional characterization of related genes.

## Materials & Methods

### Plant materials and RNA extraction

The residual heterozygous recombinant lines (RHLs) were screened from the RIL population. A pair of near isogenic lines (NILs) was developed from a F_8:9_ recombine inbred line N0910-81 by self-crossing and named RHL81-L and RHL81-S according to the difference between thousand-grain weight (TGW) and grain length (GL). Compared with RHL81-S (TGW: 33.35 g; GL: 5.36 mm), RHL81-L with a larger TGW (52.58 g) and GL (7.68 mm). The pair of NILs in this study includes four small-effect QTLs (*QTGW.nwafu-5DL.1*, *QTGW.nwafu-5DL.2*, *QTGW.nwafu-7AS.1* and *QTGW.nwafu-7AS.2*). Plants were grown in the experimental field in College of Agronomy of Northwest A&F University during the 2019-2020 growing season and grains were tagged at pollen mergence stage. Grains were removed from the spikes in the field at 4, 7, and 10 days post-anthesis stages (DPA), immediately frozen in liquid nitrogen and stored at −80°C. Total RNA was extracted using TRIZOL reagent and cDNA synthesis was performed with a reverse transcription reagent kit according to the manufacturers’ protocols.

### Identification and sequence analysis of wheat invertase genes

In order to identify the TaINV proteins, genome-wide data for Chinese Spring wheat were downloaded from the IWGSC (International Wheat Genome Sequencing Consortium) (http://www.wheatgenome.org/) to construct a local database. Two methods, blastP and HMMsearch, were used to identify gene family members initially. In detail, a wheat-specific HMM file was constructed by the HMMER build tool using the hidden markov model (HMM) seed files of invertase family (PF0000251, PF08244, PF11867, and PF12899) which downloaded from the Pfam database (http://pfam.xfam.org/) and used to screen the putative INV proteins using the HMMER search tool. Meanwhile, the protein sequences of 17 INVs in arabidopsis and 18 INVs in rice obtained from Ensemble plants database (http://plants.ensembl.org/index.html) were used as queries to search against the local protein database using an e-value of 1e-10 and an identity of 75% as the threshold. Finally, all candidate invertase obtained from the above methods were identified by CD-search (https://www.ncbi.nlm.nih.gov/Structure/cdd/wrpsb.cgi) and Pfam database.

Molecular properties about these INV genes, including the amino acid lengths, molecular weight (MW) and isoelectric point (pI) was analyzed using the ExPASy online tools (https://web.expasy.org/cgi-bin/compute_pi/pi_tool) (Artimo et al. 2012). Furthermore, the subcellular location was predicted with the SignalP 3.0 server (http://www.cbs.dtu.dk/services/SignalP/), TMHMM server (http://www.cbs.dtu.dk/services/TMHMM/), ProtComp 9.0 (http://www.softberry.com/) and WoLF PSORT (https://wolfpsort.hgc.jp/) web tools.

### Phylogenetic tree construction and gene duplication analysis

To obtain the evolutionary relationships of the invertase gene family, the 214 putative protein sequences of three monocotyledonous species, *Triticum aestivum*, *Oryza stativa*, *Zea may* and two dicotyledonous species, *Arabidopsis thaliana* and *Glycine max*, were used to construct the phylogenetic tree using the Maximum-like method in MEGA X (Kumar et al. 2018). All positions contained gaps and missing data were eliminated. The percentage of replicate trees in which the associated taxa clustered together in the bootstrap test (1000 replicates) is shown next to the branches. The display of the phylogenetic tree was optimized by using the Interactive Tree Of Life (iTOL) online website (https://itol.embl.de/upload.cgi).

### Conserved motif, gene structure and cis-acting elements

In order to identify the structural divergence of TaINV, conserved motifs in the encoded proteins were analyzed using the MEME (Multiple Expectation Maximization for Motif Elicitation) online program (http://meme-suite.org/tools/meme, version 4.12.0). Parameters were set as follows: distribution of motif occurrences - 0 or 1 per sequence, the maximum number of motifs - 10; minimum motif width - 6; and maximum motif width - 20; all other parameters were default. The sequence 2000 bp upstream of the start codon was considered the promoter region for each gene. The coding sequence and promoter sequence of *TaINV* genes were extracted from the wheat 2.1 genome. Putative functional promoter elements were analyzed using the PlantCARE (https://bioinformatics.psb.ugent.be/webtools/plantcare/html/) (Magali Lescot & Yves Van de Peer 2002). All these results, together with the intron-exon distribution, were visualized with TBtools (Chen et al. 2020).

### Chromosome localization and syntenic analysis

The physical position of all *TaINV* genes obtained from the gff3 files of wheat databases was displayed using TBtools. Gene duplication events between wheat and its relatives comprising (*Triticum urartu*, *Triticum dicoccoides*, and *Aegilops tauschii*), rice, maize, arabidopsis and soybean, were analyzed using the MCScanX (Multiple Collinearity Scan toolkit: http://chibba.pgml.uga.edu/mcscan2/) with a threshold e-value <1e-5 and 5 hits. And a collinearity map was drawn by Dual Systeny Plotter software in TBtools. Further, the substitution rate of nonsynonymous (Ka) and synonymous (Ks) was calculated by the TBtools software to determine whether selective pressure is applied to duplication events. For estimation of the timing of duplication events formula T = Ks/λ × 10^-6^ Mya was used to calculate divergence time (T) in millions of years (Mya), where λ=6.5 × 10^-9^ represented the rate of replacement of each locus per herb plant year (Peng et al., 2012; Quraishi et al., 2011).

### Expression patterns of wheat invertase gene by RNA-seq Datasets

In order to understand the functions of the *TaINV* genes, the multiple transcriptome data of different tissues (root, spike, grain, leaf, and stem) of wheat cultivar Chinese Spring and common stress (drought, heat, low temperature, *fusarium graminearum* (*FHB*), Blumeria graminis f. sp. Tritici (*Bgt*) and *Puccinia striiformis f. sp. Tritici* (*Pst*)) were downloaded from the Wheat Expression Browser (http://www.wheat-expression.com/)(Borrill et al. 2016). The spatio-temporal expression of *TaINV* genes were visualized by TBtools based on log2 (TPM + 1) values. For convenience, the heat map of genes in response to stress was drawn based on the log2 (TPM + 1) ratios of treatment to control groups. Furthermore, in order to understand the role of *TaINV* gene in grain development more clearly, the TPM values of 126 *TaINV* genes in the near-isogenic lines RHL81-L and RHL81-S was extracted from RNA-seq data (the detailed RNA-seq process is unpublished). TBtools visualized the heat map of *TaINV* genes based on log2 (TPM + 1) values.

### Quantitative real-time PCR

Nine *TaINV* genes were analyzed by a SYBR Green-based real-time quantitative PCR (qRT-PCR) with cDNA prepared from samples collected at 4, 7, and 10 DPA. The samples of RHL81-S at the same time points were set as the controls. And three independent biological replicates were prepared for each time points. Specific primers of *TaINV* and *TaActin* genes (i.e., internal reference) were designed with the Primer 5 program and shown in Table S4. qRT-PCR was performed on the QuantStudio 7 Flex Real-Time PCR System (Life Technologies Corporation) with Quantitect SYBR I Green (TaKaRa Biotechnology [Dalian] Co.Ltd) according to the manufacturer’s protocol. The 2^-ΔΔCT^ method was used to calculate the relative expression level. All data were analyzed and evaluated by GraphPad Prism 5 software. One-way analysis of variance with t-tests was performed to distinguish the differences between mean values on the same date. A value of p<0.05 was considered to be statistically significant.

## Results

### Identification and analysis of wheat INV proteins

By entering the protein sequence of the invertase identified in arabidopsis and rice, 4 HMM seed files were obtained: Glyco_hydro_32C (PF08244), Glyco_hydro_32N (PF00251), Domain of unknown function (PF11867) and Glyco_hydro_100 (PF12899). Then 192 protein sequences obtained through the HMMsearch program and BlastP were submitted to the Pfam and CDD databases for identification. Finally, 61 TaCWI, 44 TaVI, and 21 TaCI were designated TaCWI1~61, TaVI1~44 and TaCI1~21, following the nomenclature that was proposed on the chromosome designation (Table S1). The CDS lengths of the 126 *TaINV* ranged from 1371 to 2271 bp (Table S2), deduced invertase proteins contain 456-756 amino acid residues (Table S3), and their MW range from 51.37 to 84.88 kDa, similar to the invertase in arabidopsis and rice (Table S1). In addition, the putative invertase proteins encoded by TaVI, TaCWI and TaCI genes hold pI ranging from 4.77 to 6.91, 4.69-9.28, and 5.35-8.24, respectively (Table S1). Based on all the subcellular locations prediction results, we speculate that 12 TaCWIs may be targeted and located outside the cell by binding to the cell membrane, and the remaining TaCWI are secreted proteins that are located outside the cell. Most TaVIs have a signal anchor, characteristic of single-pass membrane proteins destined for lysosomal compartments. In detail, there are 3 type III-membrane protein, 8 Type I-membrane protein, 31 Type II-membrane proteins and 2 non-secretory proteins in the VIN group, suggesting that most of TaVIs are sorted to the vacuole in a N_in_/C_out_ (N-terminus in the cytoplasm, C-terminus out of the cytoplasm) membrane-bound form. Most of the TaCIs were considered as non-secretory proteins located in cytoplasmic. At the same time, TaCI1, TaCI3-5, TaCI8, TaCI10, TaCI13 and TaCI15 have a signal peptide, and they may perform signal functions by producing glucose as a substrate of hexokinase in chloroplasts or mitochondria (Table S1).

### Phylogenetic analysis of the invertase family in wheat

To determine the evolutionary relationships and reveal the subfamily classification of the INV family, a phylogenetic tree was constructed based on the alignment of all the putative full-length amino acid sequences of 214 proteins (Table S3). The phylogenetic tree showed that the TaINVs have a more closely phylogenetic relationship related to monocotyledon (Oryza stativa, Zea mays) in each clade when compared with all plants (Figure 3). The 214 INV proteins from these five species could be classified into three major groups with high confidence: CWI, VI, and CI. In the CI and VI group, the given genes from monocotyledonous and dicotyledonous plants were clustered on the same branch, confirming that CI and VI were formed before the differentiation of monocotyledonous and dicotyledonous plants. And in the VI group, genes homologous to *OsVIN2* have undergone a large number of replication events in wheat, resulting in 41 genes other than *TaVI1*, *TaVI2 and TaVI3*. While in the CWI group, the genes were clearly clustered into two subgroups, among which the α group contained all dicotyledonous genes and some monocotyledonous genes, and the β group only contained monocotyledonous genes. This indicates that the CWI gene existed before the monocotyledonous differentiation. After the monocotyledonous differentiation, the CWI gene was amplified in a large amount in the monocotyledonous plants.

**Figure 1.**
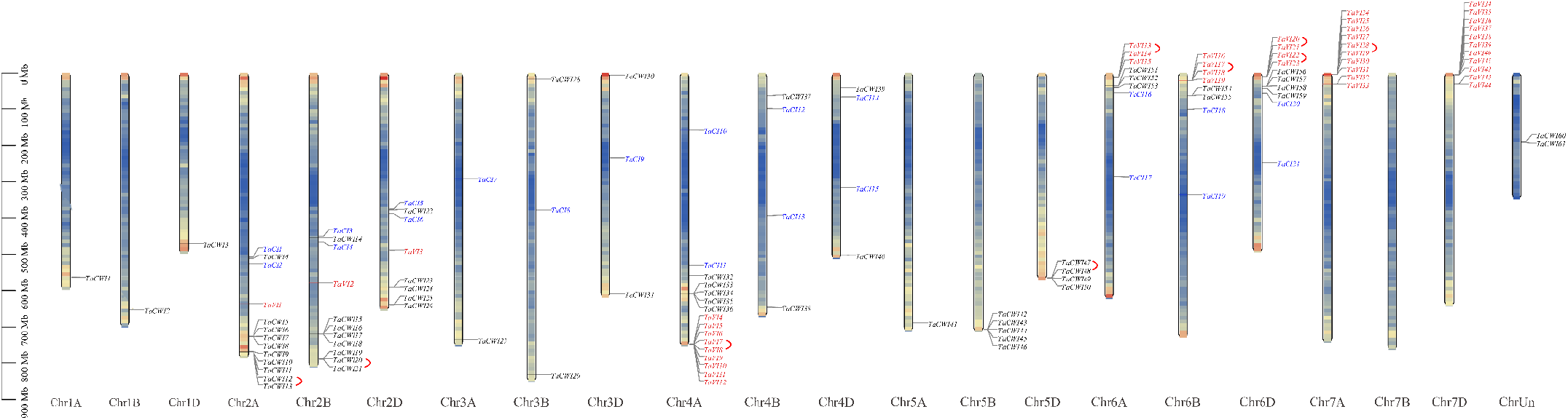
Chromosomal localization of the TaINVs. Different groups of TaINVs are represented in different colors. Black represents CWIN group, red represents VIN group and blue represents CIN group. In addition, tandem repeat genes are connected with red brackets.

**Figure 2.**
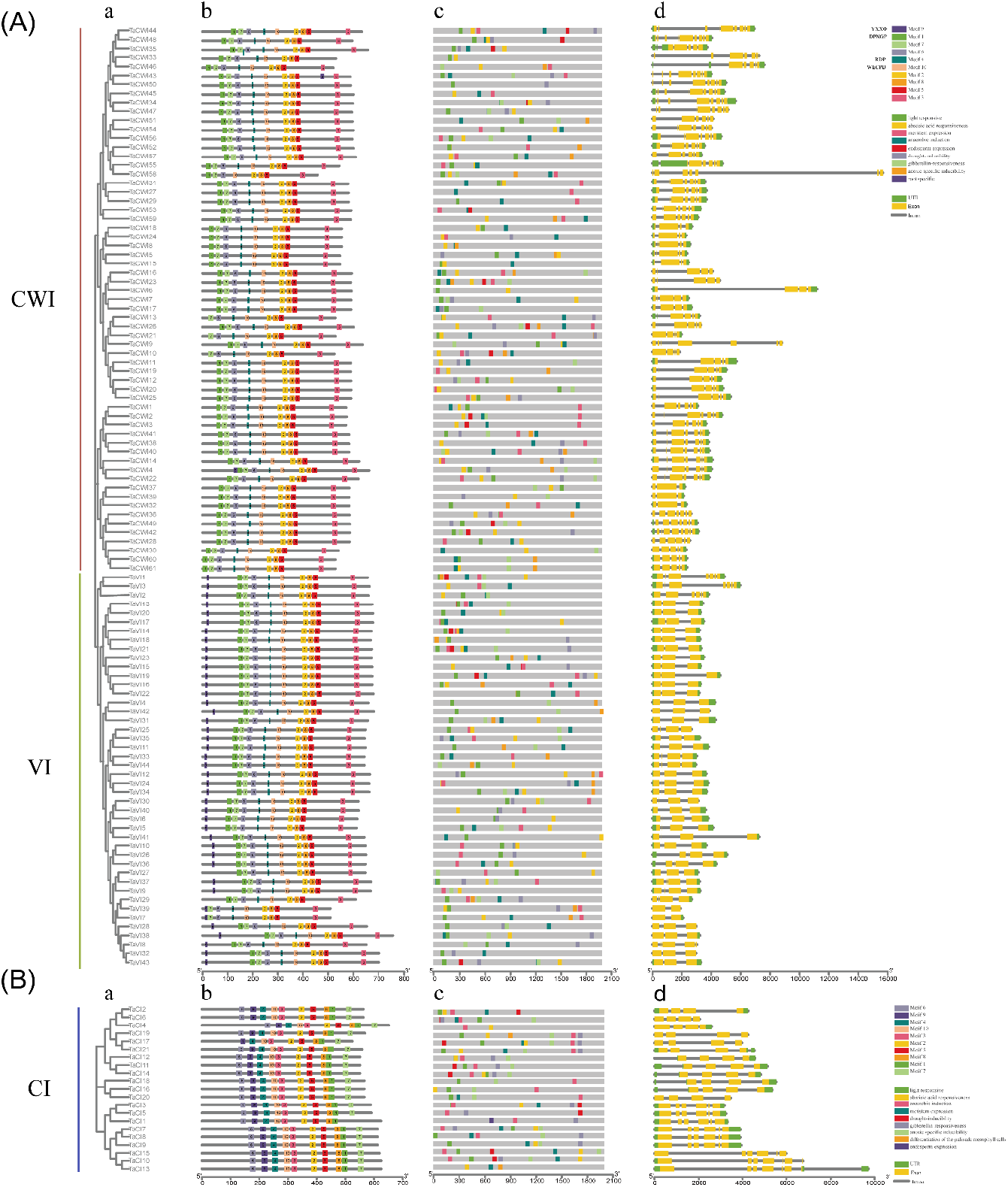
Phylogenetic relationship, motif distribution, cis-acting elements and gene structure analysis of acid invertase (A) and neutral/alkaline invertases (B). (a) Wheat TaINVs were classifed into three groups according to bootstrap values and the phylogenetic analysis of wheat; (b) ten conserved motifs identifed in protein sequences of TaINVs; (c) cis-acting elements distribution and (d) gene structures of *TaINVs*. Exons and introns were indicated by boxes and lines respectively.

**Figure 3.**
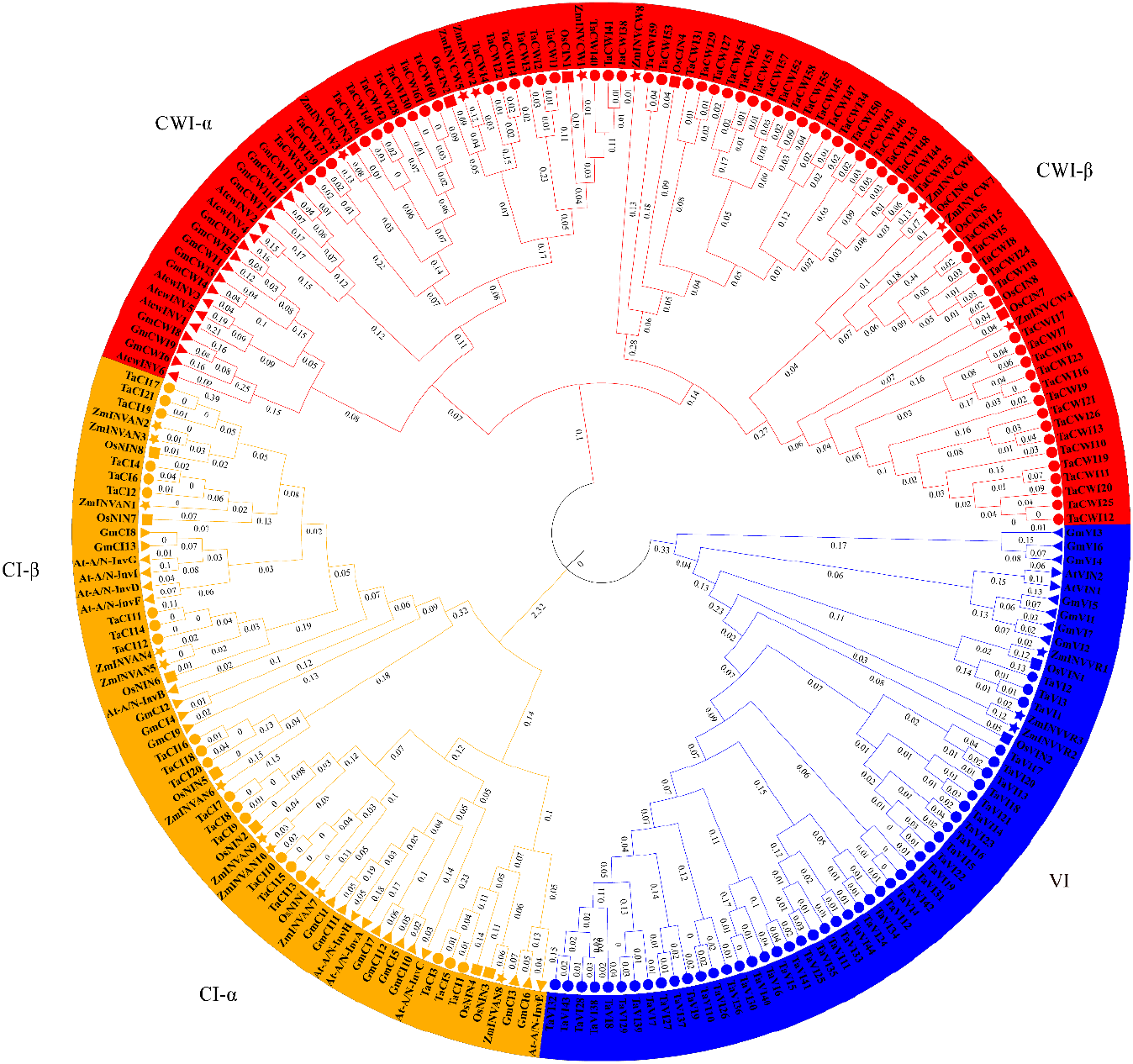
Phylogenetic analysis of the invertase family members in *Arabidopsis thaliana*, *Oryza sativa*, *Zea may*, *Glycine max*, and *Triticum aestivum*. The three main clades are distinguished by three colors, and the INVs of different species are labeled with different shapes.

### Analysis of *TaINV* Gene Structure, Motif Distribution, *Cis*-acting Elements

Analyses of gene structures showed that the exon/intron distribution patterns were similar among INV members within each clade of the phylogenetic tree, but there are some differences in the lengths of the introns among genes (Figure 2d). The exon-intron pattern of the *TaCI* gene is relatively conservative. In the CI-α group, all genes contain 6-8 exons and in the CI-β group, all genes contain 4-5 exons. It is worth noting that the genes in the α group contain a smaller exon encoding 16 amino acids. Unlike the *TaCI* gene, acid invertase has more types of exon-intron structure. Specifically, *TaVI* has 2-7 exons (without 6) and the *TaCWI* gene has 3-11 exons (without 10). The 7 and 4 exons structure genes account for the majority of *TaCWI* and *TaVI*, respectively.

To further examine the similar conserved motifs among the deduced amino acid sequences, the MEME tool was used to search for the conserved motifs in the 126 TaINVs, and 10 conserved motifs were identified (Figure 2b). In TaCI class, all genes contain the same 10 motifs with some difference in some amino acids. All acid invertases have the same 7 motifs, among which motif 2, motif 7, and motif 8 contain β-sandwich domain; motif 6 and motif 10 contain chemical substrate binding sites; motif 5 have no a clear function to date. Motif 1 (β-fructosidase motif) contains the DPN, encoded by the smallest mini-exon, which is present in most acidic invertase. A β-sandwich motif was existed in all acid invertase and had 4 conserved hydrophobic amino acid (G, A, F, and D) and 2 consecutive conserved hydrophilic amino acid (RR), which may have an important role in the correct folding of proteins. It is worth noting that a motif 9 contain the YXXØ consensus for a tyrosine-based lysosomal sorting signal and specifically exists at the N terminal of VI.

Furthermore, the *cis*-acting elements of the promoter region among 126 invertase members were analyzed and the results were shown in Figure 2c. The results showed that the *cis*-acting elements present in the promoter region of the *TaINV* genes could be divided into ten categories: light-responsive, abscisic acid responsiveness, anaerobic induction, meristem expression, gibberellin-responsiveness, drought-inducibility, anoxic specific inducibility, endosperm expression, differentiation of the palisade mesophyll cells, root-specific and other elements. Among them, the *cis*-acting elements related to light-responsive were particularly abundant, and there are 124 genes have light-responsive elements except for *TaCWI3* and *TaCWI53*. The hormone response elements are the most, with 117 and 68 genes related to hormone ABA and GA, respectively. In addition, 21 genes contain an endosperm-specific expression of *cis*-acting elements, 70 genes contain CCAAT-box, which is the MYBHv1 binding site and only *TaCWI31* has root-specific *cis*-acting elements.

### Chromosome localization and syntenic analysis of *TaINV* genes

Except for *TaCWI60* and *TaCWI61*, which are located in scaffolds, the remaining 124 genes are located on the remaining 20 chromosomes except for 7B (Figure 1, Figure 4). The *TaINV* distributions among different chromosomes are imbalanced. The number of *TaINV* genes in chromosome group A is 51 at most, followed by 41 in group D and at least 32 in group B. The chromosome 4A with 16 *TaINV* genes is the highest of any chromosome. In particular, *TaCWI* class just was absent from chromosomes 7A, 7B, and 7D, of which there are at most 10 genes on 2A, and two *TaCWI* genes are not mapped on any chromosome. *TaVI* class is distributed in the second, sixth, and seventh homologous groups, eleven *TaVI* genes are distributed on the 7D chromosome, genes homologous to 7A and 7D are located on the 4A chromosome, which is caused by the exchange of chromosomes. While, *TaCI* class is evenly distributed in homologous groups 2, 3, 4, and 6, and except for one gene on each of the 3A, 3B and 3D chromosomes, there are two *TaCI* located on the remaining chromosomes, respectively. This uneven distribution may be attributable to differences in the size and structure of the chromosomes.

**Figure 4.**
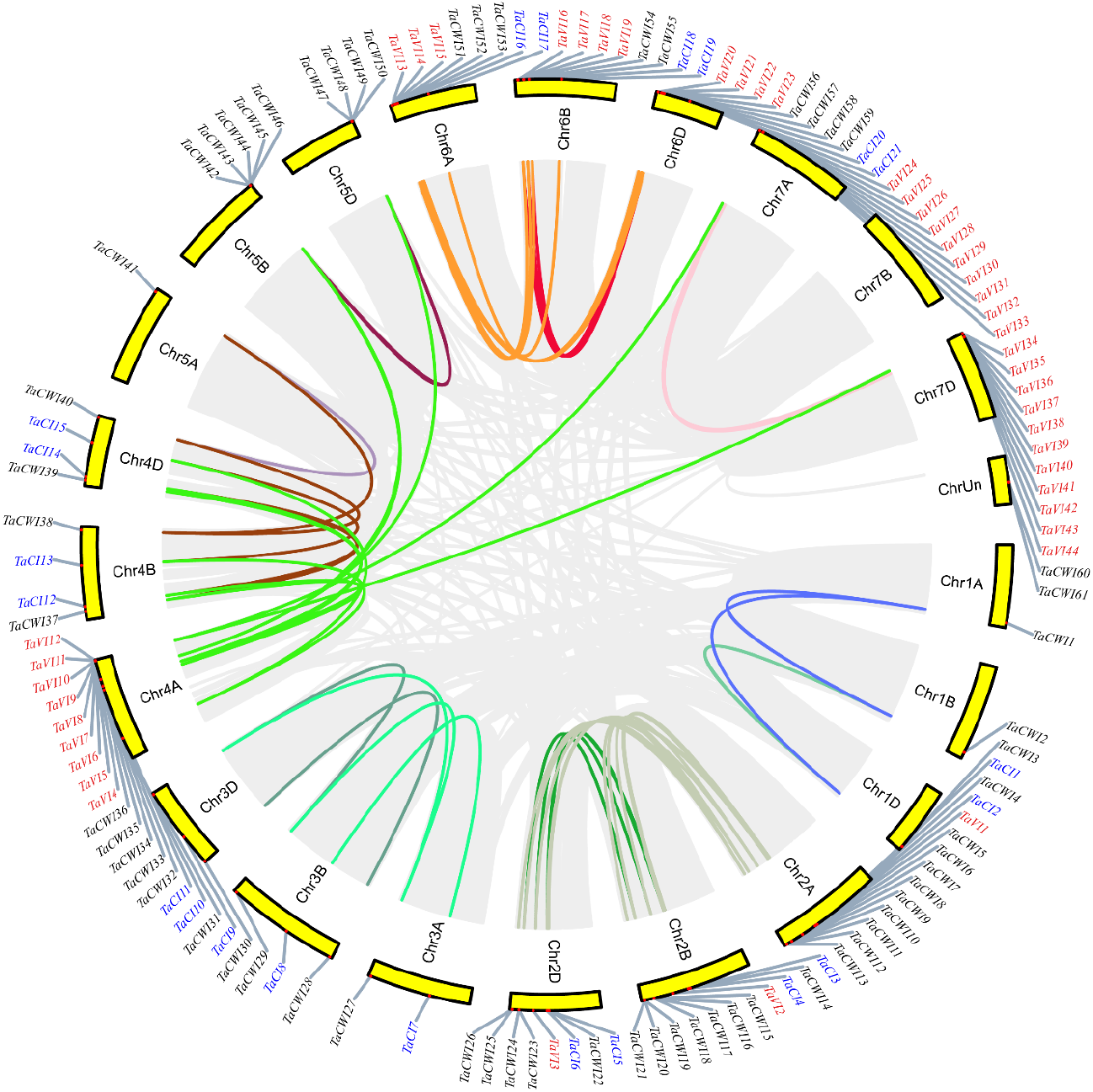
Genomic distribution of *TaINV* genes and gene homology analysis in wheat. Gray lines are all synteny blocks in the wheat genome, and the different color lines indicate duplicated *TaINV* gene pairs on different chromosome. *TaCWI*, *TaVI* and *TaCI* genes were marked by black, red and blue, respectively.

Microcollinearity analysis of the genome segments in the different species showed that the replicated segments containing the INV gene in wheat, rice and maize genomes are broadly collinear, where they form multiple sets of orthologous segments. The results also showed that 31 *TaINV* genes share homology with those in rice, 25 *TaINV* genes share homology with those in maize (Table S5). In total, 20 *TaINV* genes are directly homologous in rice and maize. There are 65 *TaINV* genes, including all 21 *TaCI*, 30 *TaCWI*, and 14 *TaVI*, that have a collinearity relationship with their ancestor species indicated that allopolyploid events were the main reason for the expansion of the *TaINV* gene family in hexaploid wheat (Table S6). In addition, the 11 and 83 gene pairs were identified as tandem and segmental duplication genes among 126 *TaINV* genes, respectively (Figure 4, Table S4). The nonsynonymous (Ka) ratio to synonymous (Ks) provided a standard for judging whether there is selective pressure on duplication events. The Ka/Ks ratios of *TaINV* duplications genes varied from 0 to 0.850892, indicating that the duplicated *TaINV* gene has undergone purification selection (Table S4).

### The expression pattern of *TaINV* genes in wheat

Gene expression patterns often contain clues to the function of the gene. The transcriptome results showed that the *TaINV* genes exhibited different temporal-spatial expression patterns (Figure 5, Table S7) and had different response patterns to stress (Figure6, Table S8). There are a total of 54 *TaINV* gene involved in stress response, including 10 *TaCI* genes, 18 *TaCWI* genes and 26 *TaVI* genes. Among them, 31 *TaINV* genes specifically respond to one of the 6 stress (dought, cold, heat, *Pst*, *Bgt* and *Fhb*), and the remaining 23 genes respond to more than 2 kinds of stress. For example, *TaCI21* is differentially expressed after *Pst* infection, while *TaCWI6*, *TaCI14*, *TaVI19*, and *TaVI22* are induced by *Bgt* pathogen. And *TaCWI30* and *TaVI31* are implicated in responding to *Fhb* infection. Intriguingly, *TaVI30* could be activated by drought, cold, heat, *Pst* and *Bgt* stress but the expression patterns showed different up-expression levels. Additionally, *TaVI35* were up-expressed in response to drought, cold, heat stress but down-expressed in response to powdery mildew stress. *TaCWI40* is up-regulated by stripe rust and fusarium graminearum, and down-regulated by cold stress. These differential genes provide a cue for the research of plant resistance to biotic and abiotic stresses.

**Figure 5.**
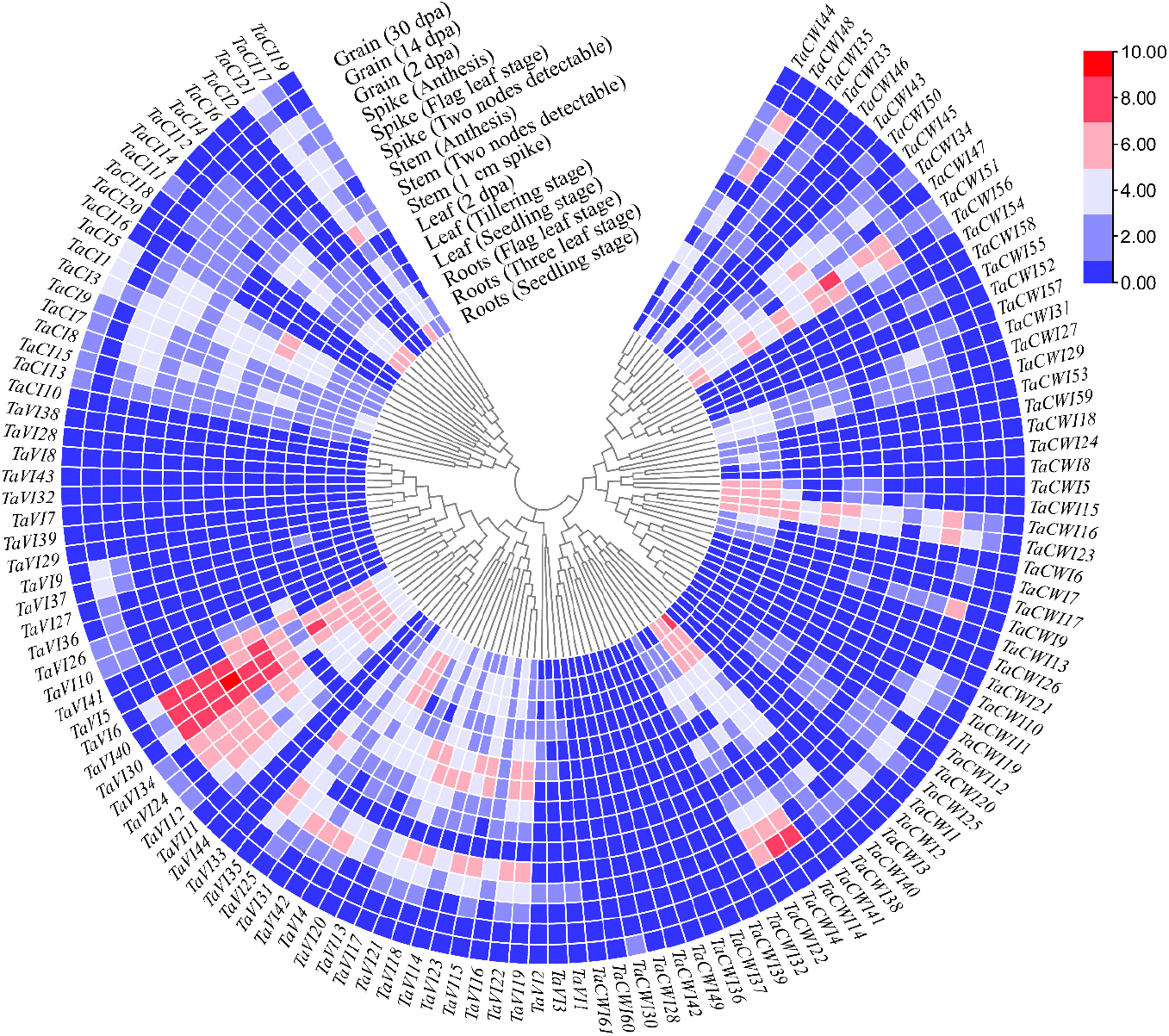
The expression profiles of *TaINV* genes involved in 5 tissues at different growth stages. The color scale of heatmap shows the level of gene expression.

**Figure 6.**
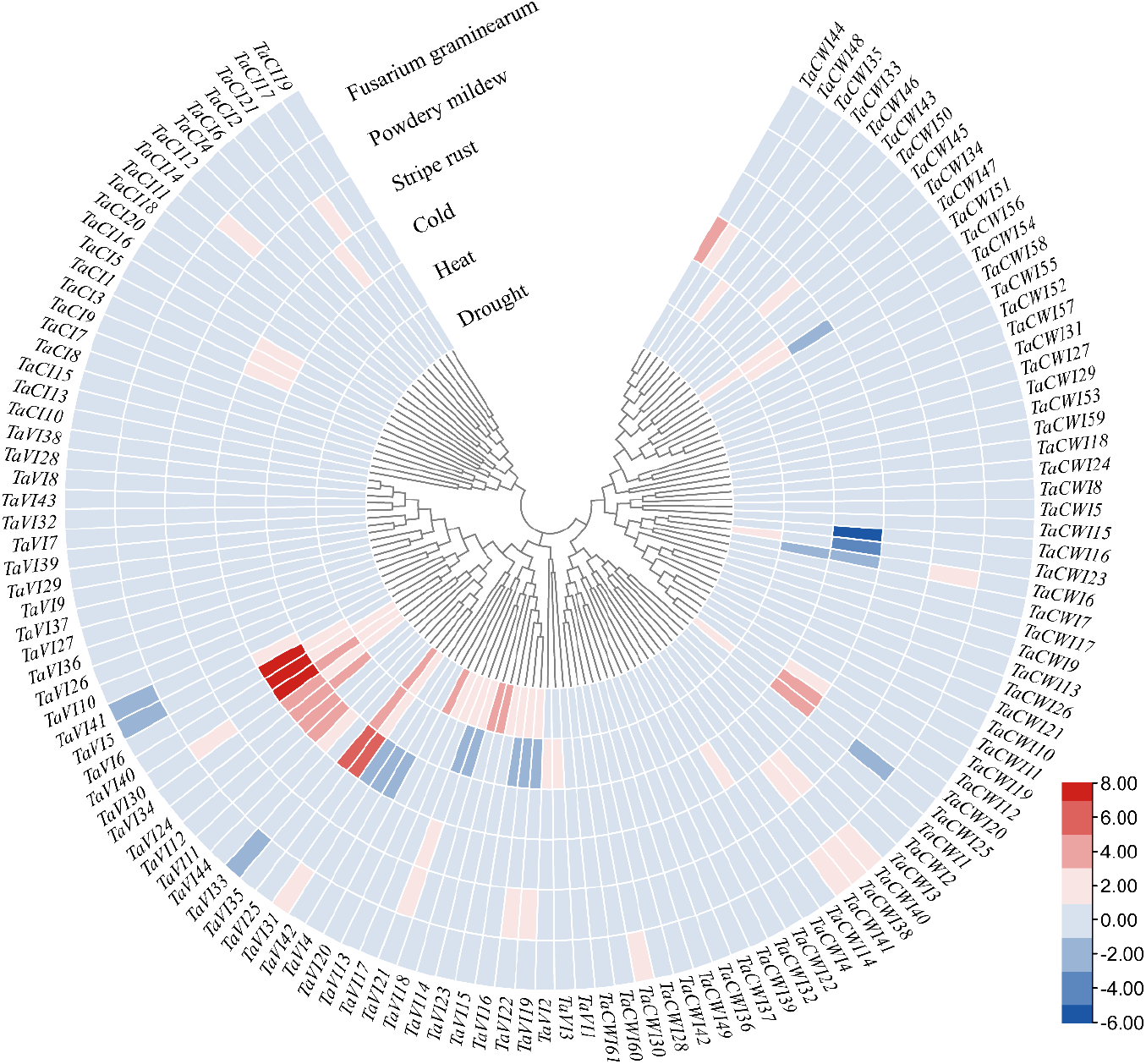
The expression profiles of TaINV genes in response to abiotic and biotic stress treatments. The expression data of 126 TaINV genes were involved in abiotic stress (drought, heat and cold) and biotic stress treatments (powdery mildew, stripe rust and fusarium graminearum) were obtained from expVIP.

As for temporal-spatial expression patterns, there have three genes (*TaCI3*, *TaCI21* and *TaCWI51*) were expressed in all tissues and developmental stages and 18 genes showed tissue-specific expression. In detail, *TaCI4* and *TaCI20* were expressed specifically in the leaf and spike, respectively. Four genes (*TaCWI17*, *TaCWI18*, *TaCWI53* and *TaCWI59*) exhibited root-specific expression and 12 genes (*TaCWI1*, *TaCWI2*, *TaCWI3*, *TaCWI10*, *TaCWI19*, *TaCWI30*, *TaCWI50*, *TaVI9*, *TaVI10*, *TaVI26*, *TaVI27* and *TaVI36*) exhibited grain-specific expression. And 29 genes were not expressed in all tissues and developmental stages, including 17, 10, 2 genes in CWIN, VIN, CIN, respectively.

### The difference in the expression of *TaINV* genes between grain weight NILs

Since a large number of genes (12 of 18) are specifically expressed in grains, we speculate that *TaINVs* may play an important role in grain development. In order to further study their roles in regulating of grain size, the TPM of *TaINVs* were obtained from a near-isogenic line after RNA-Seq. A total of 53 *TaINV* genes were detected as the different expressed genes during early grain development, comparing NILs with large grainsize and small grainsize respectively, including 19 *TaCWI*, 28 *TaVI* and 6 *TaCI* genes (Figure S1). However, only 4 genes (*TaCWI19*, *TaCWI50*, *TaVI9* and *TaVI27*) out of 12 grain-specified expressed genes exhibiting differentially expressed between near-isogenic lines (Figure S1).

Considering the limitation of the tested grain development timepoints in previous literature and the existing relationship between grain with other tissues, we selected 9 different expressed genes (not limited in Ac-INV) in RNA-Seq data to test the expression pattern in grain size NILs. The results of qRT-PCR confirmed that the grain-specific *TaINVs* had significant roles in grain development (Figure 7). At 4 DPA, except for *TaVI13* and *TaCI7*, 7 genes showed a significant difference, and the expression levels of genes (*TaVI27*, *TaCWI47*, *TaCWI50* and *TaCI17*) were higher in large grain, but *TaVI11*, *TaCWI48* and *TaCI8* were higher in small grain. At 7 DPA, the expression level of gene *TaVI13* was higher in large grain, but *TaVI11*, *TaCWI47*, *TaCI7*, and *TaCI8* were higher in small grain. At 10 DPA, except for *TaVI13*, *TaVI27*, and *TaCI17*, there are 6 genes (*TaVI11*, *TaCWI47*, *TaCWI48*, *TaCWI50*, *TaCI7*, and *TaCI8*) manifested as a significant difference and had higher expression levels under large grain. Obviously, compared with small grain, *TaCWI47* and *TaCWI50* are expressed a higher level under large grain at 4 DPA and 10 DPA of grain development.

**Figure 7.**
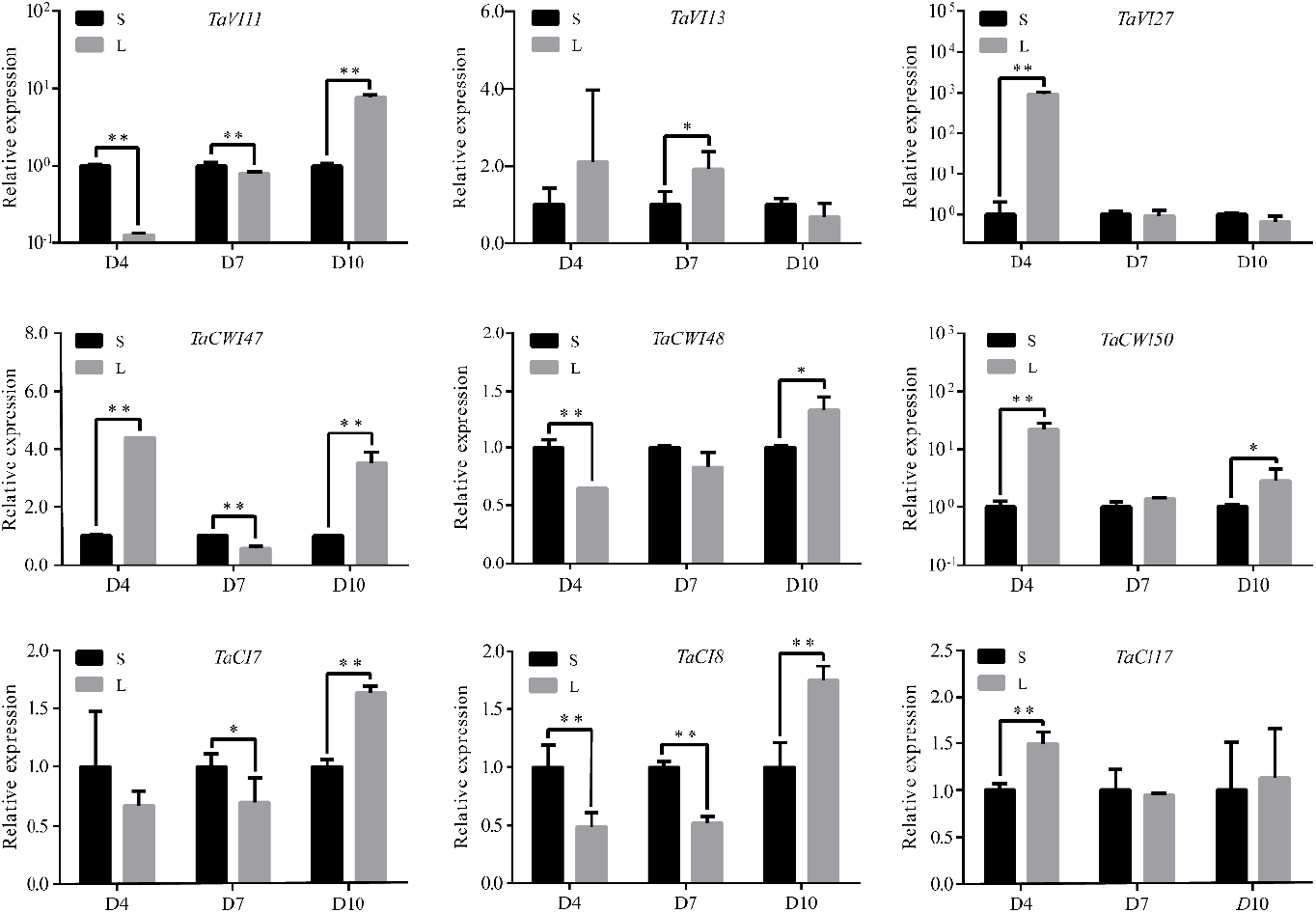
Expression profiling of 9 *TaINV* genes in grains at D4, D7 and D10 stage between RHL81-L (L) and RHL81-S (S).

## Discussion

The INV genes were widely distributed in different plant species, such as rice (Ji et al. 2005), maize (Juarez-Colunga et al. 2018), soybean (Su et al. 2018), and apple (Tae Kyung Hyun 2011) and play important roles in multiple processes of plant growth and resistance against environmental stressors (Fernández et al., 2011). However, there is no comprehensive analysis of the INV gene family in wheat (*Triticum aestivum* L.).

In this study, a total of 126 *TaINV* genes were identified from the wheat genome, containing 61 *TaCWI*, 44 *TaVI* and 21 *TaCI* genes. The *TaINV* distributions among different chromosomes are imbalanced. The chromosome 4A with 16 *TaINV* genes is the highest of any chromosome. And except for *TaCWI10* which is located in 4AS, the other 15 genes are located on chromosome 4AL. Five *TaCWI* genes from *TaCWI32* to *TaCWI36*, and 9 *TaVI* genes from *TaVI4* to *TaVI12* were located in the reciprocal translocations location between bread wheat chromosome arms 4AL and 5AL and between 4AL and 7BS, respectively. All TaCI genes are collinear (Figure 4) and have a collinearity relationship with their ancestor species (Table S6), reflecting that the amplification of *TaCI* is mainly caused by the polyploidy event of wheat species formation. There are 2 *TaCWI* genes, *TaCWI60* (*TraesCSU03G0230800.1*) and *TaCWI61* (*TraesCSU03G0235500.1*), located in scaffolds and with identical gene sequence (Table S2). Thus we speculate that *TaCWI60* and *TaCWI61* are homologous genes derived from recent replication events or actually they are the same gene.

Compared with the *TaCI*, *TaCWI* and *TaVI* genes have more exon-intron patterns (Figure 2), which are the representative traces of gene family evolution. And the lack of the mini-exon DPN amino acids was observed in 33 acid invertase genes due to the exon skip and intron retention (Figure 2). The reports showed that DPN will be lost under cold stress (Anne-Sophie Bournay, 1996). Therefore, we hypothesize that the disappearance of the mini-exon in wheat acid invertases is the result of alternative splicing events and is formed during the evolution of wheat genes in order to adapt to the cold environment.

TaCWI and TaVI have the same 9 motifs, and their position order is also the same. Compared with TaCWI, TaVI has a longer N-terminal and a unique YXXØ motif. The DPNGP, RDP and WECPD motifs each contain an acidic residue at an equivalent position in all acid invertases and it has been shown that these three motifs are indispensable for binding and catalysis (Lammens et al., 2009). In wheat acid invertase genes, the D residue in RDP motif and the E residue in WECPD motif are highly conserved in all acid invertase, while the D residue in DPNGP motif is conserved in other acid invertases except TaCWI1, TaCWI3, TaCWI10, TaCWI13, TaCWI28, TaCWI30, TaCWI36, TaCWI43, TaCWI60, TaCWI61, TaVI7, TaVI39 and TaVI42.

Further analysis revealed that 31 *TaCWI* genes were collinear with their ancestral species, 18 were generated by fragmented replication, and 6 genes produced by tandem repeat replication (Table S4, Table S6). This indicates that in addition to polyploidization, fragmentation replication also plays an important role in *TaCWI* gene amplification. In *TaVI*, 14 *TaVI* genes are collinear with their ancestral species, 19 and 3 *TaVI* genes were generated by fragmented replication and tandem repeat replication, respectively (Table S4, Table S6). Therefore, we speculate that the main reason for *TaVI* gene amplification is the fragmentation replication event after polyploidization. Thus, it could be proposed that the tandem and segmental duplication events contributed to the expansion of the acid invertase gene family in wheat.

The known functions of gene family members can be used to predict the functions of other genes on the same branch. Collinearity and phylogenetic tree analysis showed that *TaVI14*, *TaVI16* and *TaVI21* are orthologous to the *OsVIN2* gene. Tissue-specific expression analysis showed that the expression levels of these three genes were highest in the flag-leaf panicle, which was similar to the expression pattern of *OsVIN2* gene, and *OsVIN2* played an important role in yield. Therefore, we speculate that these three genes may also play an important role in wheat yield.

Similarly, *OsCIN1* is associated with early grain filling regulating in rice (Tatsuro Hirose 2002), while *OsCIN2* regulates grain shape and weight (Wang et al. 2008). Here, we noted that *TaCWI4* and *TaCWI22*, the orthologous genes of the *OsCIN1* gene, have the highest expression level in grain at the 2 DPA, which have the same expression pattern with *OsCIN1* (specifically expressed 1-4 days after flowering). Thus, we deduced that *TaCWI4* and *TaCWI22* are important for supplying a carbon source to developing filial tissues by cleaving unloaded sucrose in the apoplast. While *TaCWI14*, the orthologous genes of *OsCIN2* gene, may also was a potential domestication gene and that such a domestication-selected gene can be used for further crop improvement. This suggested that these genes may play important roles in regulating early grain filling and grain size. Although the detailed functions need to be tested in wheat, this provided a cue for dissecting the grain yield.

The tissue expression profile analyses of the *TaINV* genes showed that many *TaINV* genes were highly expressed in spike and grain, revealing that most *TaINV* genes might function in spike and grain. The expression levels of two genes in most tandem duplication gene pairs exhibited expression discrepancy, indicating that the retention of gene duplicates might be associated to processes of tissue expression divergence (Ganko et al. 2007; Huerta-Cepas et al. 2011). For instance, *TaCWI20* was highly expressed in 15 different tissues, and the expression level of *TaCWI21* was low. Under cold stress, the expression level of *TaCWI20* was induced more, indicating that *TaCWI20* might contribute more to enhance tolerance to cold stress in wheat. Furthermore, except for *TaVI7/TaVI8* and *TaVI28/TaVI29* gene pairs, the expression levels of two tandem duplication genes exhibited expression discrepancy, indicating that the retention of gene duplicates might be associated to processes of tissue expression divergence (Ganko et al. 2007; Huerta-Cepas et al. 2011). In addition, the *TaCWI12/TaCWI13* and *TaCWI20/TaCWI21* gene pairs showed significant nonfunctionalization of *TaCWI13* and *TaCWI21*, respectively. *TaCWI47/48*, *TaCWI57/58* and *TaVI11/12*, exhibited different temporal and spatial expression characteristics, while *TaVI13/14*, *TaVI17/18*, *TaVI20/21* and *TaVI22/23* showed subfunctionalization of *TaVI13*, *TaVI17*, *TaVI20 and TaVI22*, respectively.

There are 2 gene clusters located in the QTL interval, including 4 *TaCWI* genes and 8 *TaVI* genes respectively. The adjacent genes have similar expression patterns. In detail, *TaVI24* and *TaVI25* are differently expressed between the grain weight NILs and expressed after cold, heat and drought stress. *TaVI26* and *TaVI27* have grain-specific expression. *TaVI28* and *TaVI29* are not expressed in all tissues and do not respond to any stress. *TaVI30* and *TaVI31* are highly expressed in spikes. *TaCWI47* and *TaCWI48* both respond to cold stress and are differentially expressed in the NILs of DPA4, while *TaCWI49* has anther-specific expression (Jiang, 2015) and *TaCWI50* has grain-specific expression.

In this study, although our results initially indicate that these *TaINV* genes may play a role in the regulation of growth, development, the tolerance of various stresses and different expression levels in wheat postanthesis as well, much work needs to be done to resolve this regulation mechanism in the future.

## Conclusions

In this study, we comprehensively identified and characterized 126 *TaINV* genes from the wheat genome and categorized them into three classes. Gene duplication event analyses suggested that segmental duplication events contributed more than tandem duplication in the expansion of *TaINV* family. The 126 *TaINV* genes were found to be expressed in the roots, stems, leaves, spikes and grains, but some of them showed significantly different expression and/or specifically expressed patterns in the different tissues, development stages and stresses condition. The expression profiles of *TaINV* genes from RNA-seq data revealed that more half of *TaINV* genes were highly expressed in spike and grain and participated in abiotic and biotic stress responses. Consequently, these *TaINV* genes may all play a role in the regulation of growth, development, and the tolerance of various stresses in wheat as well. In a word, our results provide the basis for a deeper understanding of the biological functions of the INV gene members in wheat.

## Author Contributions

- Chao Wang conceived and designed the experiments, performed the experiments, analyzed the data, prepared figures and tables, authored or reviewed drafts of the paper.
- Guanghao Wang, Xiaojian Qu and Xiangyu Zhang performed the experiments, prepared figures and/or tables.
- Pingchuan Deng analyzed the data, prepared figures and/or tables.
- Chunhuan Chen reviewed drafts of the paper.
- Wanquan Ji conceived and designed the experiments.
- Hong Zhang conceived and designed the experiments, prepared figures and/or tables, authored or reviewed drafts of the paper.
- All authors approved the final draft.

## Acknowledgments

This work was funded by the National Key Research and Development Program of China (grant no. 2016YFD0100302), and Natural Science Foundation of Shaanxi (2021JM-090).

## Supplemental Data

Supplemental Table S1. The detailed information of 126 *TaINV* genes.

Supplemental Table S2. The coding sequences of 126 *TaINV* genes.

Supplemental Table S3. The 214 INV protein sequences used to construct the phylogenetic tree.

Supplemental Table S4. Ka/Ks ratios of *TaINV* duplication genes.

Supplemental Table S5. Syntenic relationships of *INV* genes between wheat, rice and maize.

Supplemental Table S6. Syntenic relationships of *INV* genes between wheat and its relative species.

Supplemental Table S7. The expression levels of *TaINV* genes involved in 5 tissues.

Supplemental Table S8. The expression levels of *TaINV* genes under abiotic and biotic stress treatments.

Supplemental Table S9. The lists of primers were used for qRT-PCR.

Supplemental Figure S1. Transcriptome analyses of *TaINVs* between RHL81-L and RHL81-S during the grain development.

Supplemental Figure S2. Sequence logos of wheat invertase motifs analyzed by the MEME program. The logo map of conserved sequences of ten putative motifs in acid invertases (a) and neutral/alkaline invertases (b) were identified by MEME analysis. The overall height of the stack indicates the level of sequence conservation. The height of residues within the stack indicates the relative frequency of each residue at that position.

