## supplemental file 1 for "Genome-wide identification and expression analysis of the invertase gene family in common wheat (*Triticum aestivum* L.)": Supplemental Figures.pdf

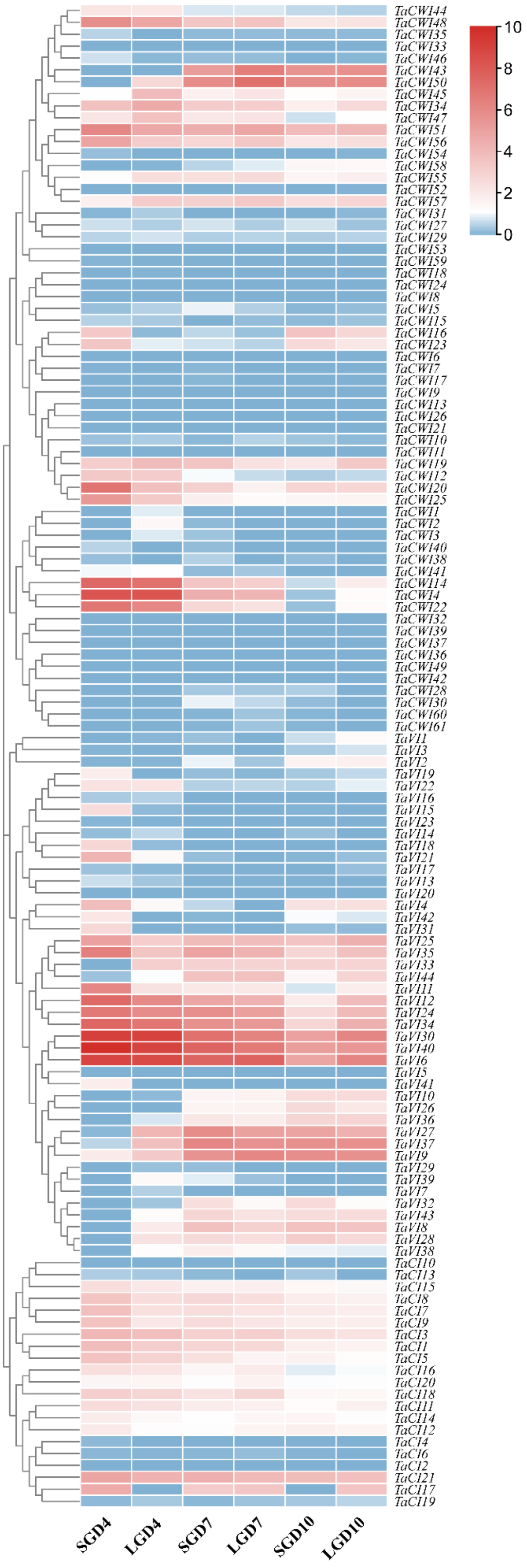


Figure S1 Transcriptome analyses of TaINVs between RHL81-L/S during the grain development.

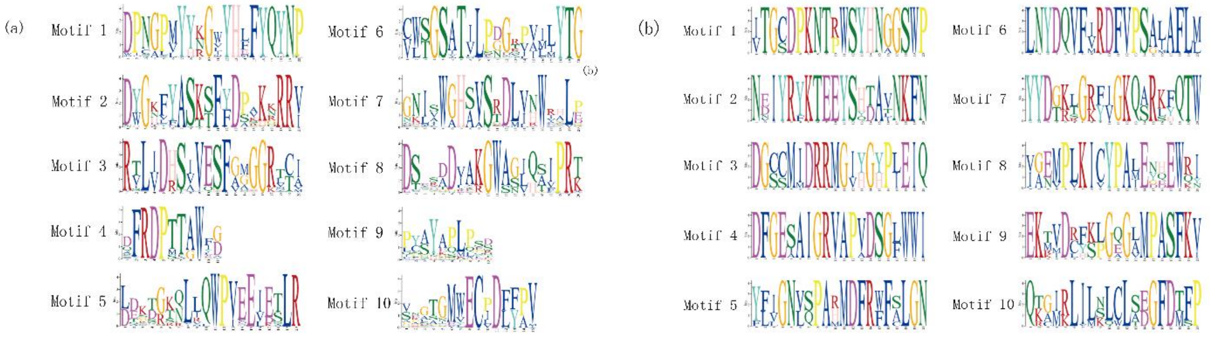

Figure S2. Sequence logos of wheat invertase motifs analyzed by the MEME program. The logo map of conserved sequences of ten putative motifs in acid invertases (a) and neutral/alkaline invertases (b) were identified by MEME analysis. The overall height of the stack indicates the level of sequence conservation. The height of residues within the stack indicates the relative frequency of each residue at that position.
